# Identification and characterization of a novel *pic* gene cluster responsible for picolinic acid degradation in *Alcaligenes faecalis* JQ135

**DOI:** 10.1101/530550

**Authors:** Jiguo Qiu, Lingling Zhao, Siqiong Xu, Qing Chen, Le Chen, Bin Liu, Qing Hong, Zhenmei Lu, Jian He

## Abstract

Picolinic acid (PA) is a natural toxic pyridine derivative. Microorganisms can degrade and utilize PA for growth. However, the full metabolic pathway and its physiological and genetic foundation remain unknown. In this study, we identified the *pic* gene cluster responsible for the complete degradation of PA from *Alcaligenes faecalis* JQ135. PA was initially 6-hydroxylated into 6-hydroxypicolinic acid (6HPA) by PA dehydrogenase (PicA). 6HPA was then 3-hydroxylated by a four-component 6HPA monooxygenase (PicB) to form 3,6-dihydroxypicolinic acid (3,6DHPA), which was then converted into 2,5-dihydroxypyridine (2,5DHP) by a decarboxylase (PicC). The 2,5DHP was further degraded into fumaric acid, through PicD (2,5DHP dioxygenase), PicE (*N*-formylmaleamic acid deformylase), PicF (maleamic acid amidohydrolase), and PicG (maleic acid isomerase). Homologous *pic* gene clusters with diverse organizations were found to be widely distributed in *α*-, *β*-, and *γ*-Proteobacteria. Our findings provide new insights into the microbial metabolism of environmental toxic pyridine derivatives.

**Importance:** Picolinic acid is a common metabolite of L-tryptophan and some aromatic compounds and is an important intermediate of industrial concern. Although the microbial degradation/detoxification of picolinic acid has been studied for over 50 years, the underlying molecular mechanisms are still unknown. Here, we show the *pic* gene cluster responsible for the complete degradation of picolinic acid into the tricarboxylic acid cycle. This gene cluster was found to be widespread in other *α*-, *β*-, and *γ*-Proteobacteria. These findings provide new perspective for understanding the mechanisms of picolinic acid biodegradation in bacteria.

## Introduction

Picolinic acid (PA) is a natural toxic pyridine derivate. PA is a dead-end metabolite of L-tryptophan via the kynurenine pathway in humans and other mammals (1, 2). It has been identified in various biological mediums, such as cell culture supernatants, blood serum, and human milk (3). PA can also be produced in other biological processes, including the microbial degradation of 2-aminophenol, nitrobenzene, catechol, anthranilic acid, and 3-hydroxyanthranilic acid (4-7) (Fig. S1). In industrial applications, PA is an important intermediate in the organic synthesis of pharmaceuticals (e.g. carbocaine), herbicides (e.g. picloram and diquat), and fungicide (e.g. 3-trifluoromethyl picolinic acid) (8, 9). Moreover, due to its chelating properties, PA is added to chromium and iron preparations as a way to treat diabetes and anemia (10, 11). Nevertheless, the toxic or disadvantages to the use of PA have also been found. PA has been shown to inhibit the growth of normal rat kidney cells and T cell proliferation, enhance seizure activity in mice, and induce cell death via apoptosis (12-14). In particular, PA showed high antimicrobial activity at concentrations as low as 8 μg/L, which is enough to inhibit the growth of Gram-positive bacteria (15-17).

No reports showed that PA could be metabolized by humans (18). Nevertheless, numerous microorganisms were found to degrade PA, such as *Alcaligenes* (19), *Arthrobacter* (20), *Burkholderia* (21), *Streptomyces* (22), and an unidentified Gram-negative bacterium (designated as UGN strain) (23). The bacterial degradation pathway of PA was proposed as: PA, 6-hydroxypicolinic acid (6HPA),3,6-dihydroxypicolinic acid (3,6DHPA), and 2,5-dihydroxypyridine (2,5DHP), according to several previous determinations of degradation intermediates and crude enzymes (Fig. 1). However, little is known about the degradation mechanism at genetic level.

**Fig. 1.**
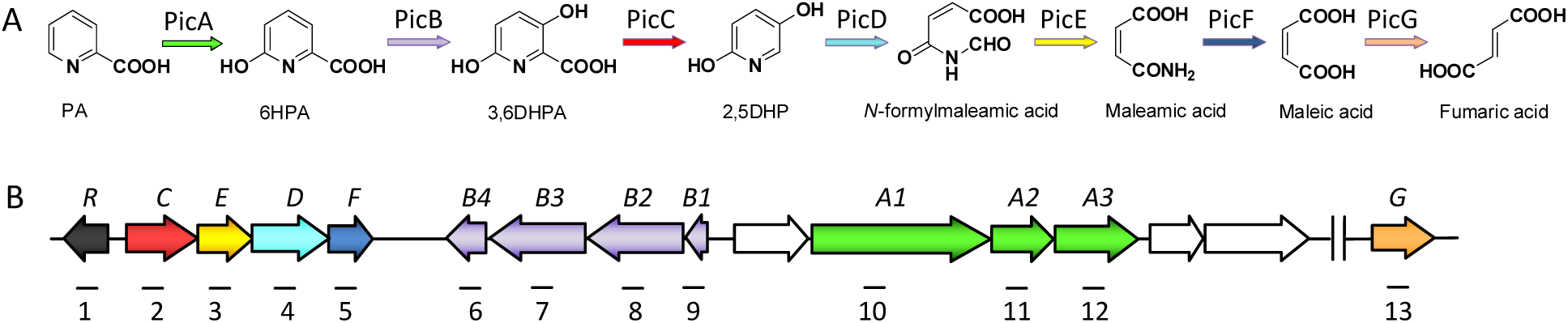
PA degradation in *A. faecalis* JQ135. (A) The complete catabolic pathway and corresponding enzymes. (B) Genetic organization of the *pic* gene cluster. Genes are annotated following the color scheme in A.

*Alcaligenes faecalis* strain JQ135 can degrade PA and utilize it for cell growth (24) (25). Our previous study demonstrated the first genetic evidence involving PA degradation, in which *picC* (encoding 3,6-dihydroxypicolinic acid decarboxylase) gene was responsible for the conversion of 3,6DHPA to 2,5DHP in *A. faecalis* strain JQ135 (26). Nevertheless, the complete degradation mechanism of PA are still unknown. In this study, we reported the *pic* gene cluster responsible for PA degradation in strain JQ135 (Fig. 1). The enzymes encoded by the *pic* gene cluster were expressed and functionally characterized using *in vitro* enzymatic assays, while the *pic* clusters in other bacteria were also predicted.

## Results and discussion

### Identification of *pic* gene cluster for PA degradation

Previous results showed that gene *AFA_15145* (*picC*) was responsible for the conversion of 3,6DHPA to 2,5DHP in *A. faecalis* JQ135 (26). Bioinformatic analysis indicated that *picC* was located in the gene cluster (designated *pic*). (Fig. 1). The *pic* cluster consisted of 13 genes and their predicted functions are summarized in Table 1. RT-PCR analysis showed that all of the *pic* genes were induced by PA, except for citrate in MSM (Fig. 2). The above results suggest that the *pic* cluster was involved in PA metabolism in *A. faecalis* JQ135.

**Table 1.**
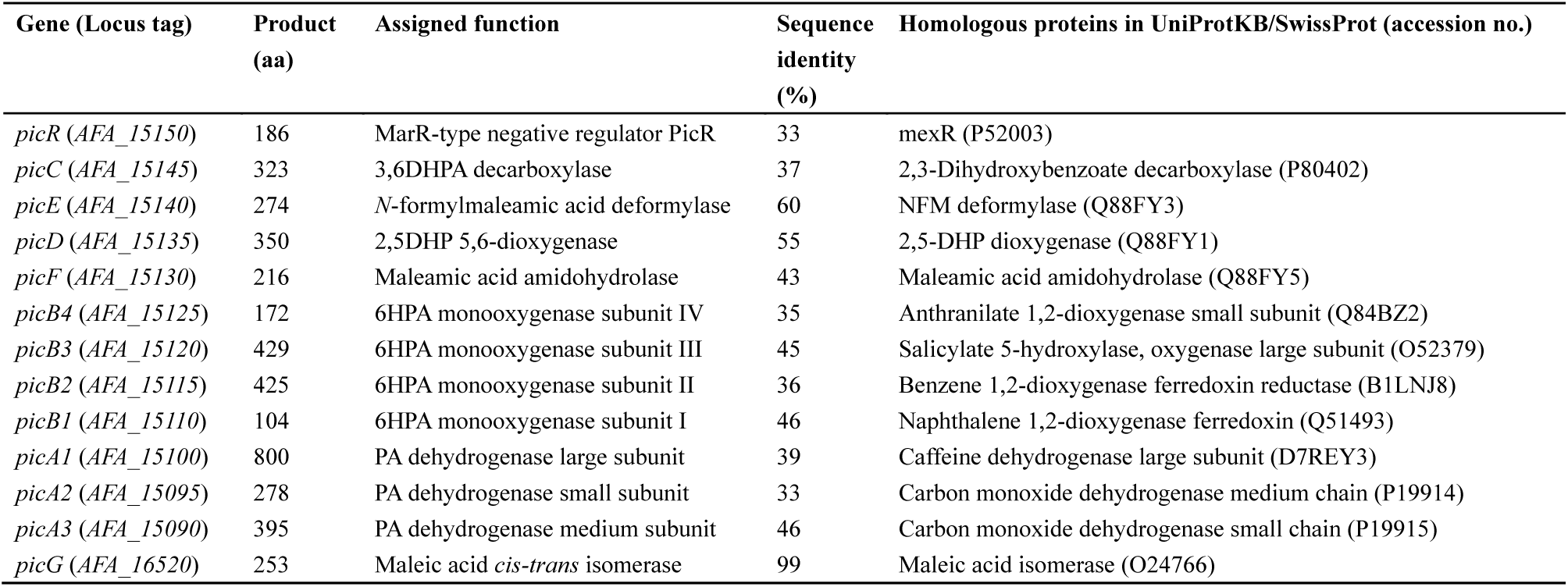
The *pic* genes and assigned function in *Alcaligenes faecalis* JQ135.

**Fig. 2.**
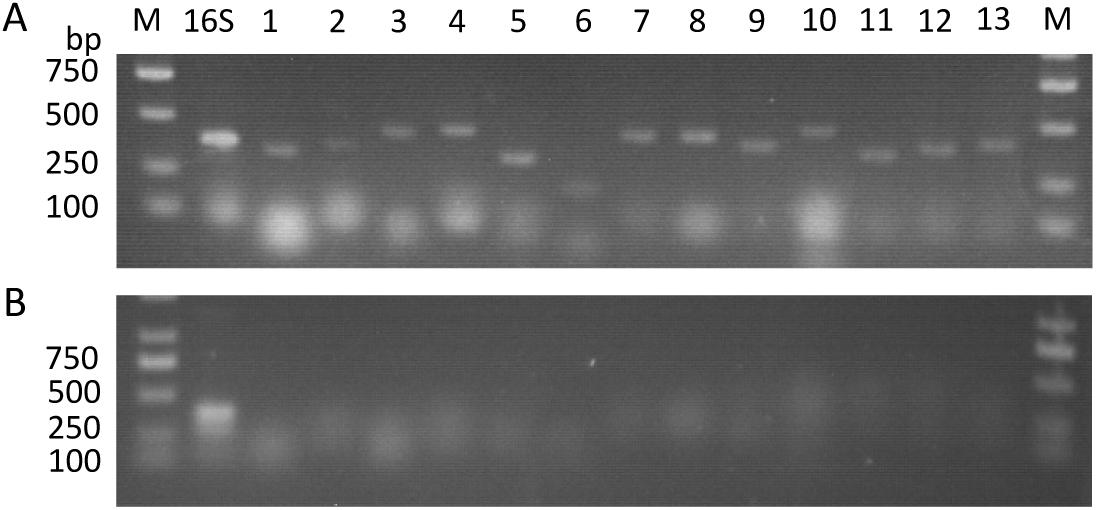
Agarose gel electrophoresis of RT-PCR products generated using RNA of *A. faecalis* JQ135. when cells grow with PA (A) or citrate (B). M, DNA marker. 16S, positive control. 1 to 13 were corresponding to the positions in Fig. 1B.

### The *picA1A2A3* encodes the PA dehydrogenase (PicA)

Previous studies reported that PA was initially 6-hydroxylated to 6HPA (19, 23, 24, 27). The hydroxylation of *N*-heterocyclic aromatic compounds were usually catalyzed by multicomponent molybdopterin-containing dehydrogenase (28, 29). In the *pic* gene cluster, the *picA1, picA2*, and *picA3* gene products showed the highest similarities (∼40%) to the respective subunits of molybdopterin-containing dehydrogenases, such as carbon monoxide dehydrogenase and caffeine dehydrogenase (Table 1). PicA1, PicA2, and PicA3 were predicted to contain binding domains for the molybdopterin cytosine dinucleotide (MCD), FAD, and two [Fe-S] clusters, respectively (Fig. 3). Thus, the *picA1A2A3* genes were considered to catalyze the first step of PA degradation. When the *picA1A2A3* genes were deleted, the acquired mutants JQ135Δ*picA1A2A3* lost the ability to grow on PA but could still grow on 6HPA (Fig. 4). The complementation strain JQ135Δ*picA1A2A3*/pBBR-*picA1A2A3* restored the phenotype of growth on PA. Moreover, the *picA1A2A3* genes were then cloned into pBBR1MCS-5 (pBB-*picA1A2A3*) and transferred into the PA-non-degrading strain *Pseudomonas putida* KT2440. The recombinant KT/pBBR-*picA1A2A3* was found to acquire the ability to convert PA into 6HPA. When lacking one component, the recombinants KT/pBBR-*picA2A3* and KT/pBBR-*picA1A2* did not show the transformation ability, indicating that all of these three components were essential.

**Fig. 3.**
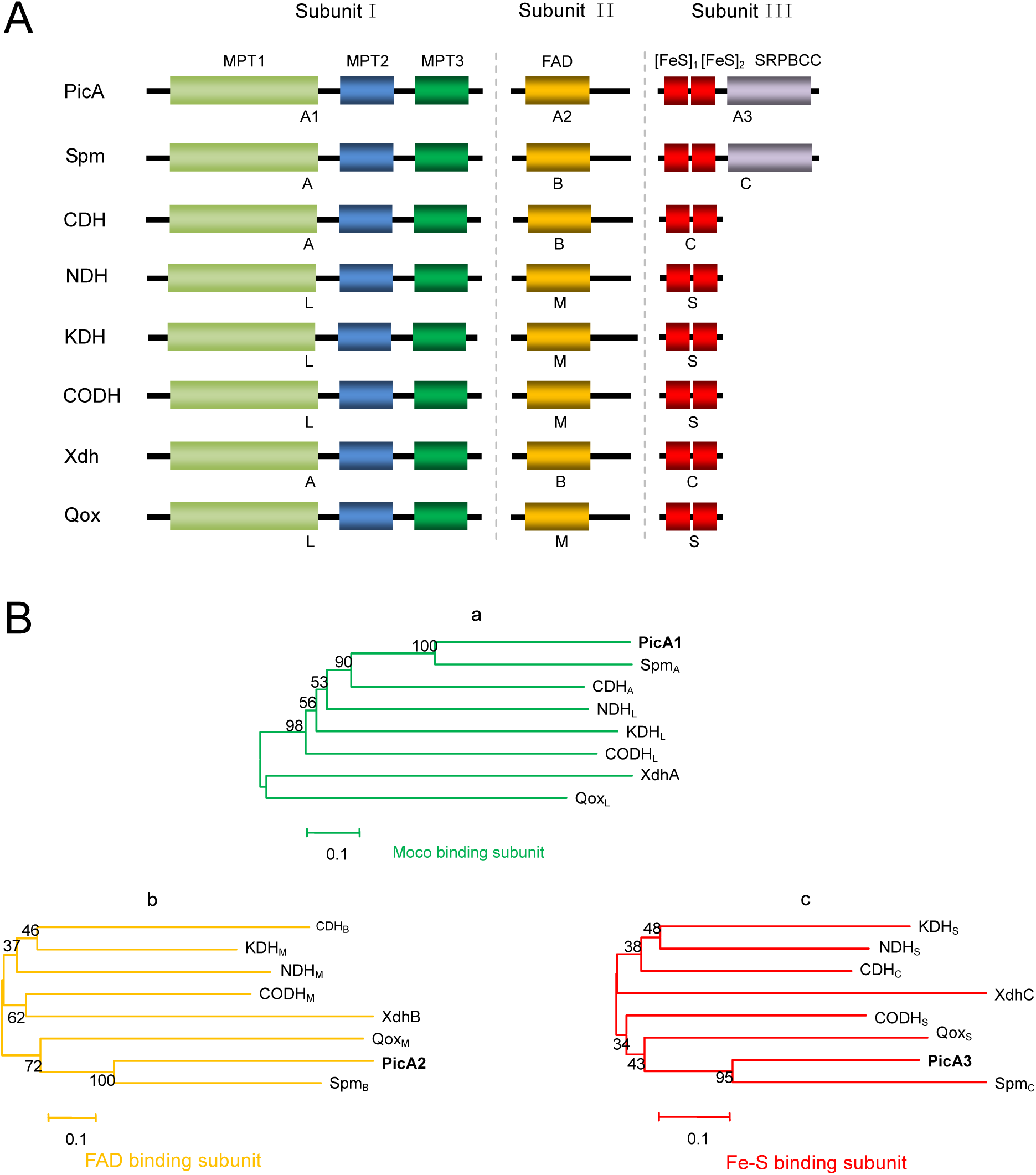
Bioinformatic analysis of PicA. (**A**) Molecular architecture of several multicomponent molybdenum-containing hydroxylases. Subunits I, II, and III are molybdopterin cytosine dinucleotide (MCD), FAD, and two [Fe-S] clusters containing components, respectively. Spm_ABC_ (GenBank accession numbers AEJ14617 and AEJ14616), 3-succinoylpyridine dehydrogenase from *P. putida*; CDH_ABC_ (D7REY3, D7REY4, and D7REY5), caffeine dehydrogenase from *Pseudomonas* sp. strain CBB1; NDH_LMS_ (CAA53088, CAA53087, and CAA53086), nicotine dehydrogenase from *Athrobacter nicotinovorans*; KDH_LMS_ (WP_016359451, WP_016359456, and WP_016359457), ketone dehydrogenase from *A. nicotinovorans*; CODH_LMS_ (P19913, P19914, and P19915), carbon monoxide dehydrogenase from *Hydrogenophaga pseudoflava*; XDH_ABC_ (Q46799, Q46800, and Q46801), xanthine dehydrogenase from *E. coli*; Qox_LMS_ (CAD61045, CAD61046, and CAD61047), quinaldine 4-oxidase from *A. ilicis*. The letters depicted below the proteins indicate the subunit names of the corresponding proteins. The conserved domains are as follows: MPT, domains for binding to the molybdopterin cytosine dinucleotide cofactor (MoCo); FAD, FAD-binding domain; [FeS], ferredoxin-like [2Fe-2S]-binding domain; SRPBCC, SRPBCC ligand-binding domain. (**B**) Phylogenetic analysis of PicA and related molybdenum containing hydroxylases. (a) Phylogenetic analysis of PicA1 and the Moco binding subunit of other enzymes.(b) Phylogenetic analysis of PicA2 and the FAD binding subunit of other enzymes. (c) Phylogenetic analysis of PicA3 and the [2Fe-2S] binding subunit of other enzymes. The phylogenetic trees were constructed by using the neighbor-joining method (with a bootstrap of 1000) with software MEGA 6.0. The bar represents amino acid substitutions per site.

**Fig. 4.**
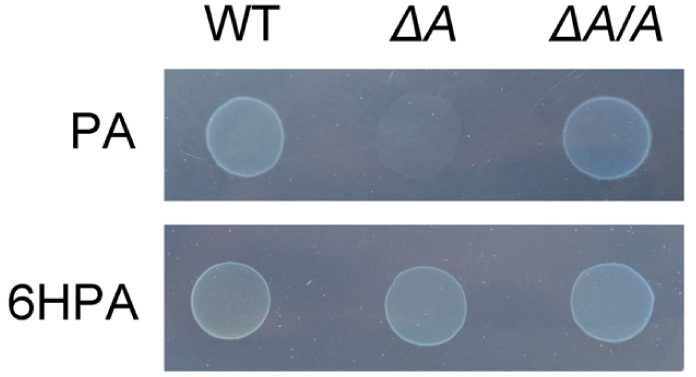
Growth phenotype of wild type *A. faecalis* JQ135 (WT), mutant JQ135Δ*picA1A2A3* (*ΔA*), and the complemented strain JQ135*ΔpicA1A2A3*/pBBR-*picA1A2A3* (*ΔA/A*) on MSM plates with 1.0 mM PA and 6HPA as sole carbon sources.

After being pre-grown in MSM containing 1.0 mM citrate and 1.0 mM PA for 12h, the cell-free extract of KT/pBBR-*picA1A2A3* produced PA dehydrogenase activity when phenazine methosulfate (PMS) was used as an electron acceptor. The *K*_*m*_ value for PA at pH 7.0 and 25°C was 0.65 ± 0.14 μM, the *v*_*max*_ was 44.89 ± 2.45 mU/mg (Fig. S2). No PA dehydrogenase activity was detected under anaerobic conditions, which was similar to the nicotinate hydroxylase (NicAB) using O_2_ as the final electron acceptor in *P. putida* KT2440 (28). All of these results demonstrated that *picA1A2A3* genes are responsible for the hydroxylation of PA into 6HPA in *A. faecalis* JQ135.

### The *picB1B2B3B4* encodes the four-component 6HPA monooxygenase (PicB)

The second step in the PA metabolic pathway is predicted to be the 3-hydroxylation of 6HPA to 3,6DHPA (23). A BLAST homology search against the database sequences revealed that four genes in the same orientation: *picB1, picB2, picB3*, and *picB4*, showed high identities (35%∼45%) at amino acid level with the respective components of bacterial Rieske non-heme iron aromatic ring-hydroxylating oxygenases (RHO) (Table 1, Fig. 5). RHOs are usually involved in hydroxylation of aromatic compounds (30, 31). Therefore, the *picB1B2B3B4* genes were most likely responsible for the degradation of 6HPA to 3,6DHPA. The *picB1B2B3B4* genes were deleted from the genome of *A. faecalis* JQ135. The mutant JQ135Δ*picB1B2B3B4* lost the ability to grow on PA or 6HPA, but could still grow on 3,6DHPA (Fig. 6). The complementation strain JQ135Δ*picB1B2B3B4*/pBBR-*picB1B2B3B4* could utilize 6HPA for cell growth. These four genes were cloned into pBBR1MCS-5 and transferred into *P. putida* KT2440. The recombinant KT/pBBR-*picB1B2B3B4* was found to convert 6HPA into 3,6DHPA. Moreover, the recombinants KT/pBBR-*picB2B3B4* and KT/pBBR-*picB1B2B3* did not exhibit transformation ability, suggesting that all of these four components were necessary. These results indicated that *picB1B2B3B4* genes are responsible for the conversion of 6HPA into 3,6DHPA in *A. faecalis* JQ135.

**Fig. 5.**
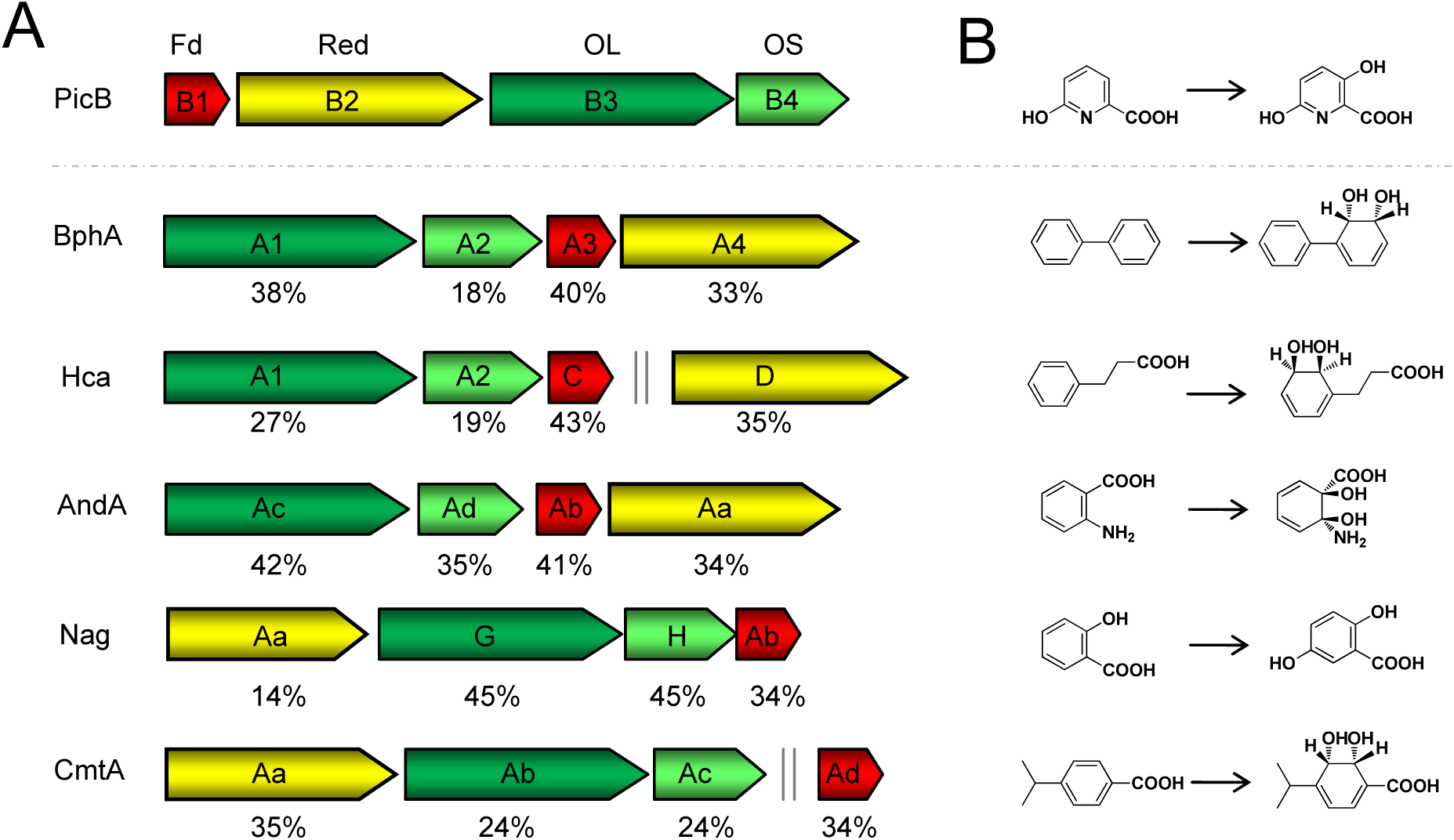
Genetic organizations of the PicB encoding genes as compared with similar Rieske non-heme iron aromatic ring-hydroxylating oxygenase genes (A) and the corresponding enzyme reactions (B). The arrows in A indicate the size and direction of each gene. Homologous genes are shown in the same color. Fd, ferredoxin; Red, reductase; OL, oxygenase large component; OS, oxygenase small component. Double vertical gray lines indicate discontinuous genes. Numbers below the arrows indicate the percent amino acid sequence identity with the ortholog *picB* gene product. BphA1A2A3A4 (GenBank accession numbers Q53121, Q53122, Q53123, and Q0S032), biphenyl 2,3-dioxygenase from *Rhodococcus jostii*. HcaA1A2CD (P0ABR5, Q47140, P0ABW0, and P77650), 3-phenylpropionate dioxygenase from *E. coli*. AndAaAbAcAd (Q84BZ0, Q84BZ1, Q84BZ3, and Q84BZ2), anthranilate 1,2-dioxygenase from *Burkholderia cepacia*. NagAaGHAb (O52379, O52380, and O52381), salicylate 5-hydroxylase from *Ralstonia* sp. CmtAaAbAcAd (Q51973, Q51974, Q51975, and Q51978), *p*-cumate 2,3-dioxygenase from *Pseudomonas putida*.

**Fig. 6.**
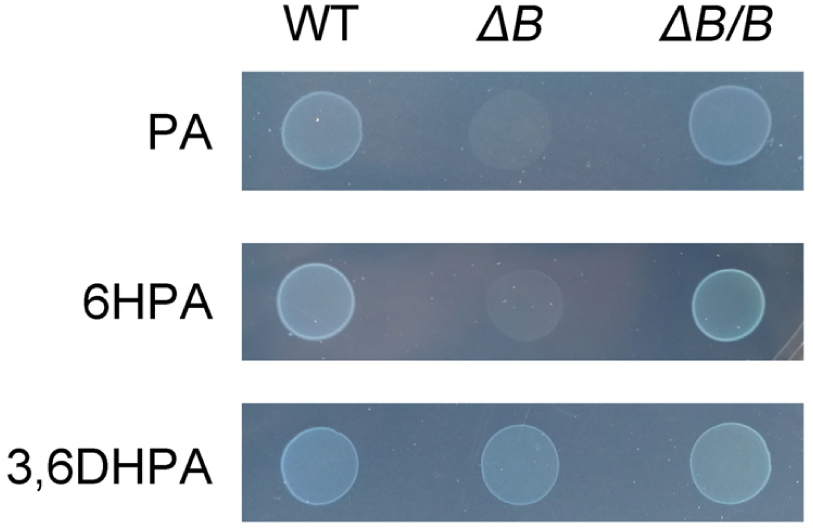
Growth phenotype of wild type *A. faecalis* JQ135 (WT), mutant JQ135Δ*picB1B2B3B4* (*ΔB*), and the complemented strain JQ135*ΔpicB1B2B3B4*/pBBR-*picB1B2B3B4* (*ΔB/B*) on MSM plates with 1.0 mM PA, 6HPA, and 3,6DHPA as sole carbon sources.

Interestingly, the cell-free extract of 6HPA-pre-grown KT/pBBR-*picB1B2B3B4* could not convert 6HPA, even though several factors (e.g. FAD, FMN, NADH, or NADPH) were added. Previous studies have attempted to purify the 6HPA monooxygenase from cell-free extract of *Arthrobacter picolinophilus* DSM 20665 (32) and the UGN strain (23), but no PicB activities were detected at all. These results suggested that the PicB was commonly fragile.

### PicDEF and PicG (MaiA) converts 2,5-DHP into fumaric acid

In *A. faecalis* JQ135, the 3,6DHPA was found to be degraded into 2,5DHP by 3,6-dihydroxypicolinic acid decarboxylase (PicC) (26). 2,5DHP is a central intermediate of numerous pyridine derivatives, including nicotinate (28), nicotine (33), 2-hydroxy-pyridine (34), and 5-hydroxypicolinic acid (25). The bacterial metabolism of 2,5DHP usually employs the maleamate pathway, whose key enzymes include 2,5DHP dioxygenase, *N*-formylmaleamate (NFM) deformylase, maleamate amidase, and maleate *cis-trans* isomerase (28, 33). The primary structure of PicD, PicE, and PicF showed high similarities (40∼60%) to 2,5DHP 5,6-dioxygenase, NFM deformylase, and maleamic acid amidohydrolase, respectively (Fig. S3), which strongly implies that 2,5DHP follows the maleamate pathway in *A. faecalis* JQ135. Then PicD was overexpressed in *E. coli* BL21 (DE3). The purified PicD showed 2,5DHP degrading ability, which was measured spectrophotometrically by a decrease in absorbance at 320 nm, with a *K*_m_ value of 65.72±6.27 μM. PicD showed no activity without 1 mM Fe^2+^, which resembles previous reports of the 2,5DHP dioxygenase from *P. putida* KT2440 and *Ocrobactrum* sp. SJY1 (28, 33). These results suggest that PicD, PicE, and PicF catalyze the formation of maleic acid from 2,5DHP in *A. faecalis* JQ135.

Maleic acid could be isomerized into fumaric acid, an intermediate of the Krebs cycle. However, the *pic* gene cluster lacks an isomerase gene. Previous study showed that *AFA_16520* (*maiA*, also named *picG* in this study) gene was only the maleic acid *cis-trans* isomerase gene in *A. faecalis* JQ135 and PicG was responsible for the last step of PA degradation (25). Furthermore, a DNA fragment containing *picEDFG* genes was cloned into a 2,5DHP non-degrading bacterium *Sphingomonas wittichii* DC-6 (35). Recombinant DC-6/pBBR-*picEDFG* acquired the ability to degrade and utilize 2,5DHP for growth (Fig. S4). While lacking the *picG* gene, recombinant DC-6/pBBR-*picEDF* was not able to grow on 2,5DHP. These results also indicated that PicDEF and PicG (MaiA) were responsible for the conversion of 2,5DHP to fumaric acid.

### Diversity of the *pic* genes in other bacteria

Six complete genome sequences from *A. faecalis* strains are available in NCBI and *pic* gene clusters were found in all of these genomes (Fig. 7, Table S1). Besides *A. faecalis*, the orthologous *pic* genes were also found in other *α-, β-*, and *γ*-Proteobacteria (21 genera and over 160 strains) (Table S1). Most of these strains belong to the order Burkholderiales of the class *β*-Proteobacteria, including genera of *Achromobacter, Alcaligenes, Advenella, Bordetella, Burkholderia, Caballeronia, Cupriavidus, Pandoraea, Paraburkholderia, Polaromonas, Pseudacidovorax, Pusillimonas, Rolstonia*, and *Variovorax*. Interestingly, some of these strains are toxic compound degraders (e.g., *Achromobacter xylosoxidans* A8 and *Variovorax* sp. JS1663) (36, 37), human pathogens (e.g., *Bordetella pertussis* Tohama I and *Bordetella bronchiseptica* RB50) (38), and nematicidal bacteria (e.g., *A. faecalis* ZD02) (39), whereas other strains represent typical soil bacteria (e.g., *Pseudomonas*), plant rhizosphere-associated bacteria (e.g., *Bradyrhizobium*), arctic bacterium (*Octadecabacter arcticus* 238) (40), and antarctic bacterium (*Hoeflea* sp. IMCC20628) (41). Previous reports also showed that bacteria in the genera of *Aerococcus, Arthrobacter*, and *Streptomyces* could degrade PA (20, 22, 27). However, no homologues of *pic* genes were found in these bacteria, suggesting that other new and unknown catabolic genes might exist.

**Fig. 7.**
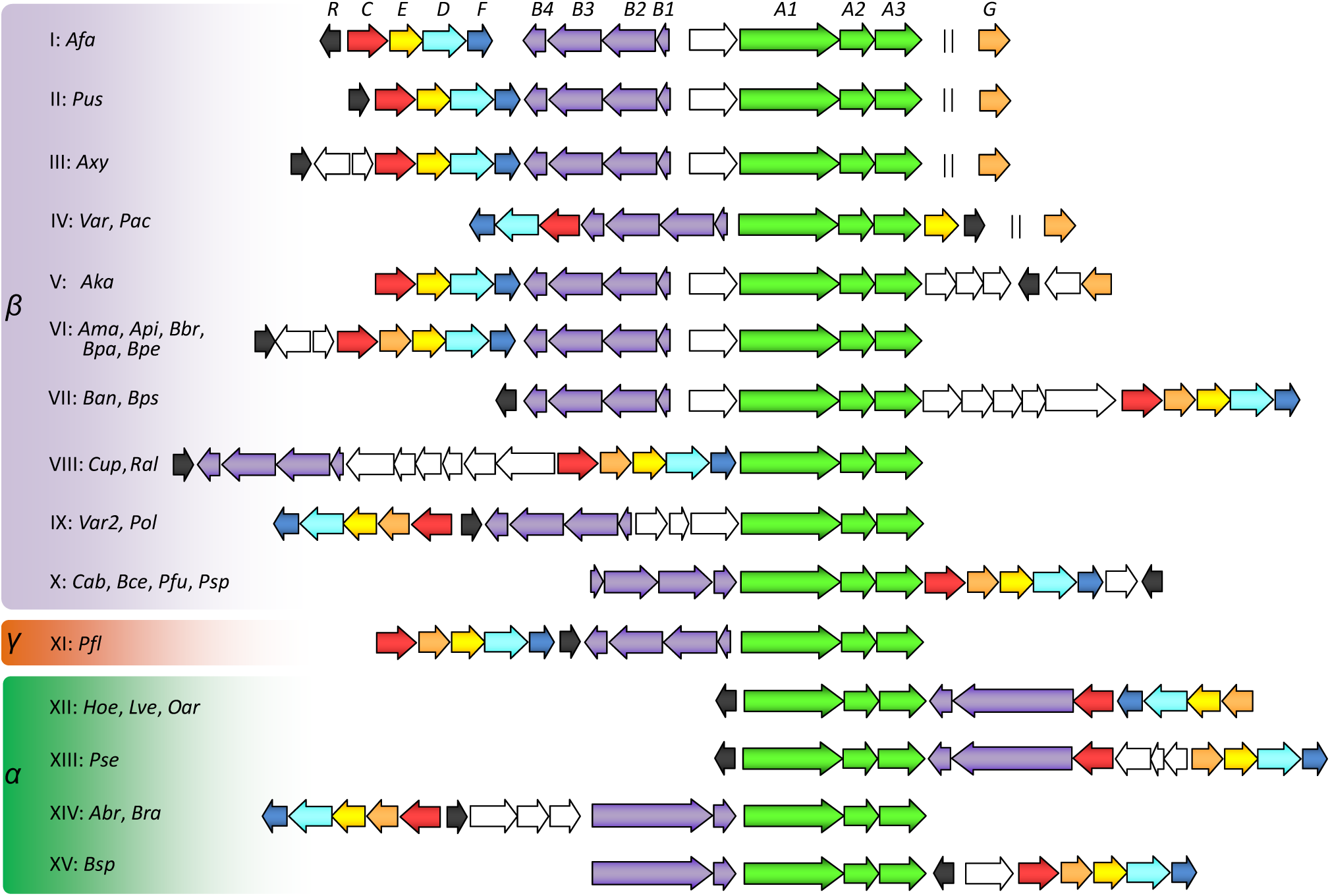
Predicted PA catabolism gene clusters in bacterial genomes. *α, β*, and *γ* indicate the *alpha-, beta-*, and *gamma*-Proteobacteria. I to XV: The 15 different types of PA catabolism loci. Abbreviations and representative strains: *Afa, Alcaligenes faecalis* JQ135; *Pus, Pusillimonas* sp. T2; *Axy, Achromobacter xylosoxidans* A8; *Var, Variovorax paradoxus* S110; *Pac, Pseudacidovorax* sp. RU35E; *Aka, Advenella kashmirensis* W13003; *Ama, Achromobacter marplatensis* B2; *Api, Achromobacter piechaudii* ATCC 43553; *Bbr, Bordetella bronchiseptica* RB50; *Bpa, Bordetella parapertussis* ATCC BAA-587; *Bpe, Bordetella pertussis* Tohama I; *Ban, Bordetella ansorpii* NCTC13364; *Bps, Bordetella pseudohinzii* HI4681; *Cup, Cupriavidus necator* N-1; *Ral, Ralstonia eutropha* JMP134; *Var2, Variovorax* sp. JS1663; *Pol, Polaromonas* sp. OV174; *Cab, Caballeronia arvi* LMG 29317; *Bce, Burkholderia cepacia* JBK9; *Pfu, Paraburkholderia fungorum* NBRC 102489; *Psp, Pandoraea sputorum* DSM 21091; *Pfl, Pseudomonas fluorescens* C8; *Hoe, Hoeflea* sp. IMCC20628; *Lve, Loktanella vestfoldensis* SMR4r; *Oar, Octadecabacter arcticus* 238; *Pse, Paracoccus seriniphilus* DSM 14827; *Abr, Afipia broomeae* GAS525; *Bra, Bradyrhizobium canariense* GAS369; *Bsp, Bradyrhizobium* sp. C9. The detailed genomic accession numbers and the gene locus tags are listed in Table S1. Identities (percent) of amino acid sequences between Pic proteins of strain *A. faecalis* JQ135 and representative homologs are listed in Table S2.

The genetic organization of the *pic* gene clusters was highly diverse (Fig. 7). The *picA* genes usually adjoin *picB* genes, except in the *Cupriavidus* strains. In most genera, the *picG* gene was located between *picC* and *picE*. However, in *Alcaligenes, Pusillimonas*, and some *Achromobacter* strains, *picG* was located at a distant site away from the *pic* gene cluster. In *β*- and *γ*-Proteobacteria, PicB consists of four components. Interestingly, in *α*-Proteobacteria, including genera of *Bradyrhizobium, Hoeflea*, and *Octadecabacter*, PicB consists of two components, in which the large subunit PicB123 encodes proteins that correspond to PicB1, PicB2, and PicB3 (Fig. S5).

### Conclusions

PA is generated through cell metabolism or synthesized artificially industry and is thus ubiquitous in the environment. The degradation and detoxification of PA by microorganisms have been studied for more than 50 years (27). This study revealed that the *pic* gene cluster is responsible for the complete degradation of PA. PicA initially converted PA into 6HPA, then PicB hydroxylated 6HPA into 3,6DHPA, which was further converted into 2,5DHP by PicC. The 2,5DHP was further degraded into fumaric acid by PicD, PicE, PicF, and PicG. The genetic functions of *picA* and *picB* genes is reported here for the first time. In addition, the findings of homologous *pic* gene cluster in other *α*-, *β*-, and *γ*-Proteobacteria would help us to understand the growth, competition, and environmental adaptability of bacteria in the natural environment.

## Methods

### Chemicals

PA, 6HPA, and 2,5DHP were purchased from J&K Scientific Ltd. (Shanghai, China). 3,6DHPA was chemically synthesized (26). Other reagents were purchased from Sangon Biotech Co., Ltd. (Shanghai, China).

### Strains, plasmids, and culture conditions

All bacterial strains and plasmids used in this study are listed in Table 2. Mineral salts medium (MSM) and Luria-Bertani medium (LB) have been described previously (25). *Alcaligenes faecalis* JQ135 was the wild-type PA-degrading strain (24). *P. putida* KT2440 is a model Gram-negative organism that cannot degrade PA (28). *Sphingomonas wittichii* DC-6 is a chloroacetanilide herbicide degrader that is not unable to degrade PA or 2,5DHP (35). *E. coli* DH5α acted as host for the construction of plasmids. *E. coli* BL21(DE3) was used to over-express proteins. Bacteria were grown in LB medium at 37 ºC (*E. coli*) or 30 ºC (*Alcaligenes, Pseudomonas, Sphingomonas*, and their derivates). Antibiotics were added as required at the following concentrations: chloramphenicol (Cm), 34 mg/L; gentamicin (Gm), 50 mg/L; kanamycin (Km), 50 mg/L; and streptomycin (Str), 50 mg/L.

**Table 2.**
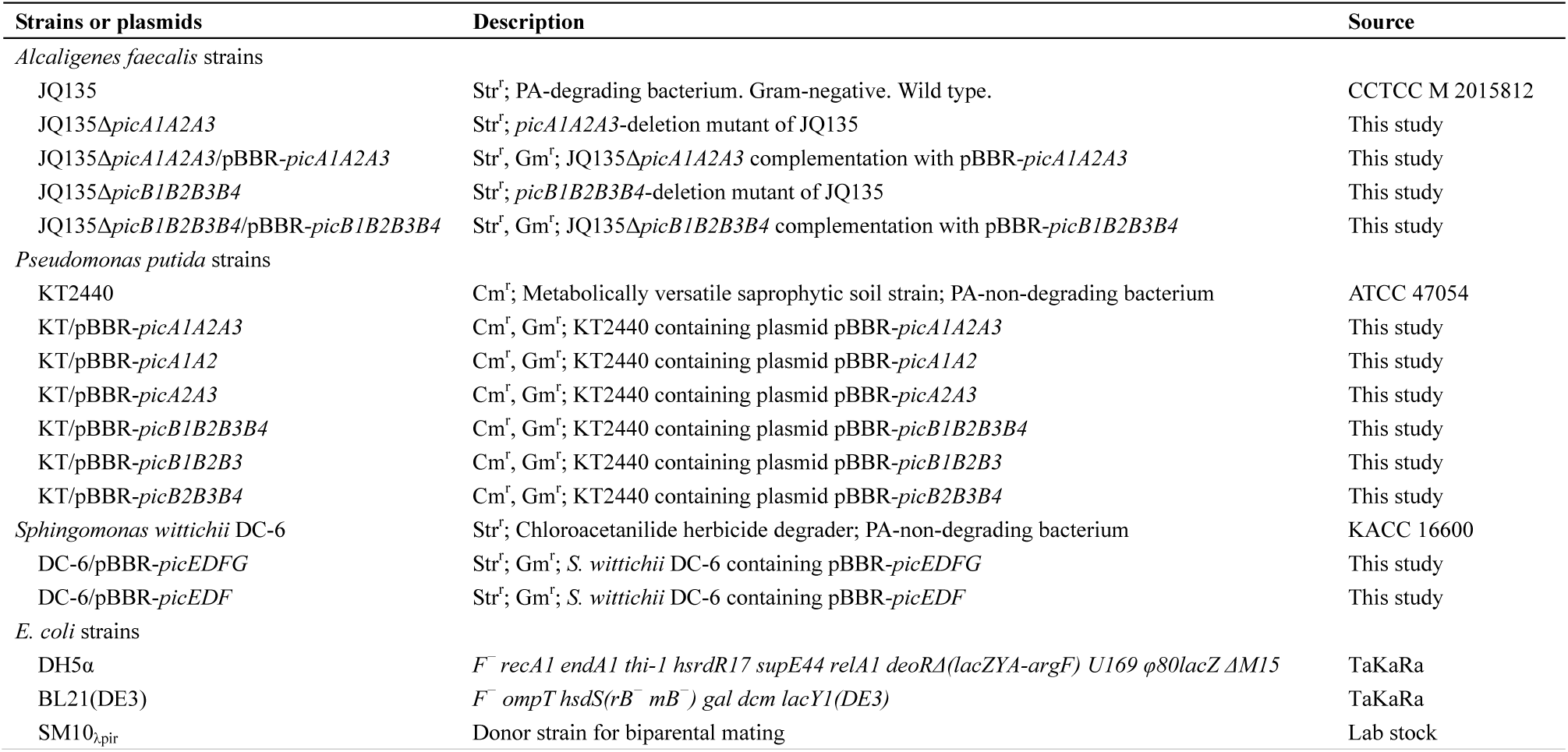
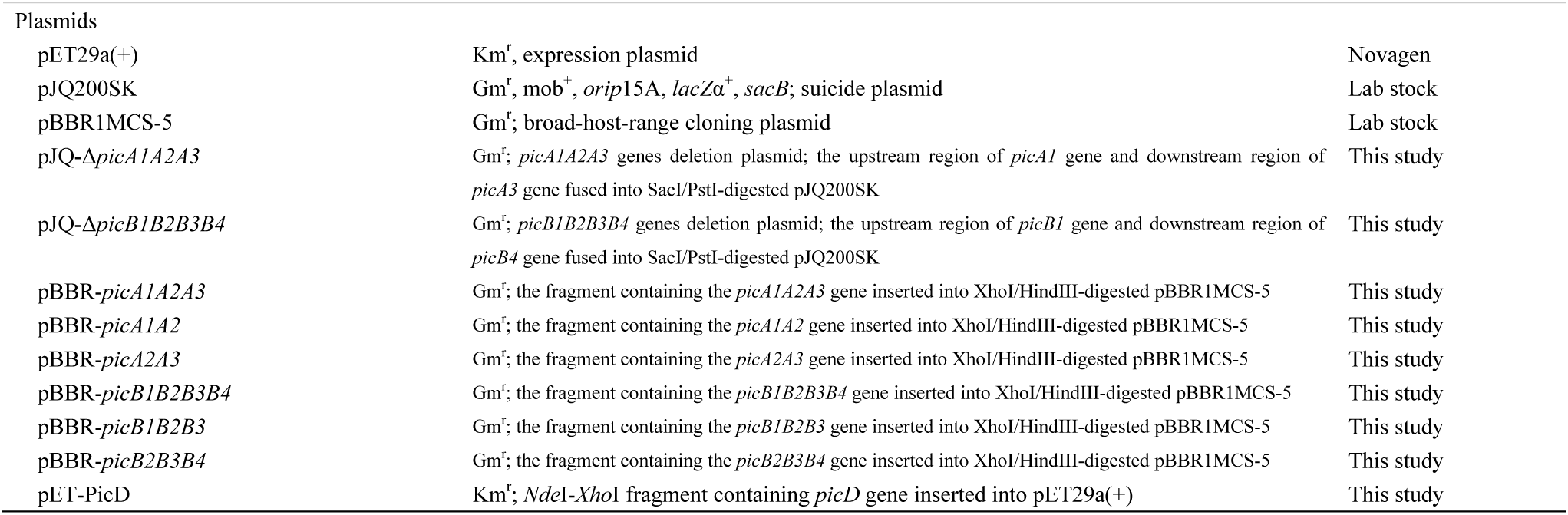
Strains used in this study.

### General DNA techniques

Routine isolation of genomic DNA, extraction of plasmids, restriction digestion, transformations, PCR, and electrophoresis were performed according to standard procedures in Molecular Cloning-A Laboratory Manual (New York: Cold Spring Harbor Laboratory). Primer synthesis and the sequencing of PCR products or plasmids were performed by Genscript Biotech (Nanjing, China). The primers used in this study are listed in Table 3.

**Table 3.**
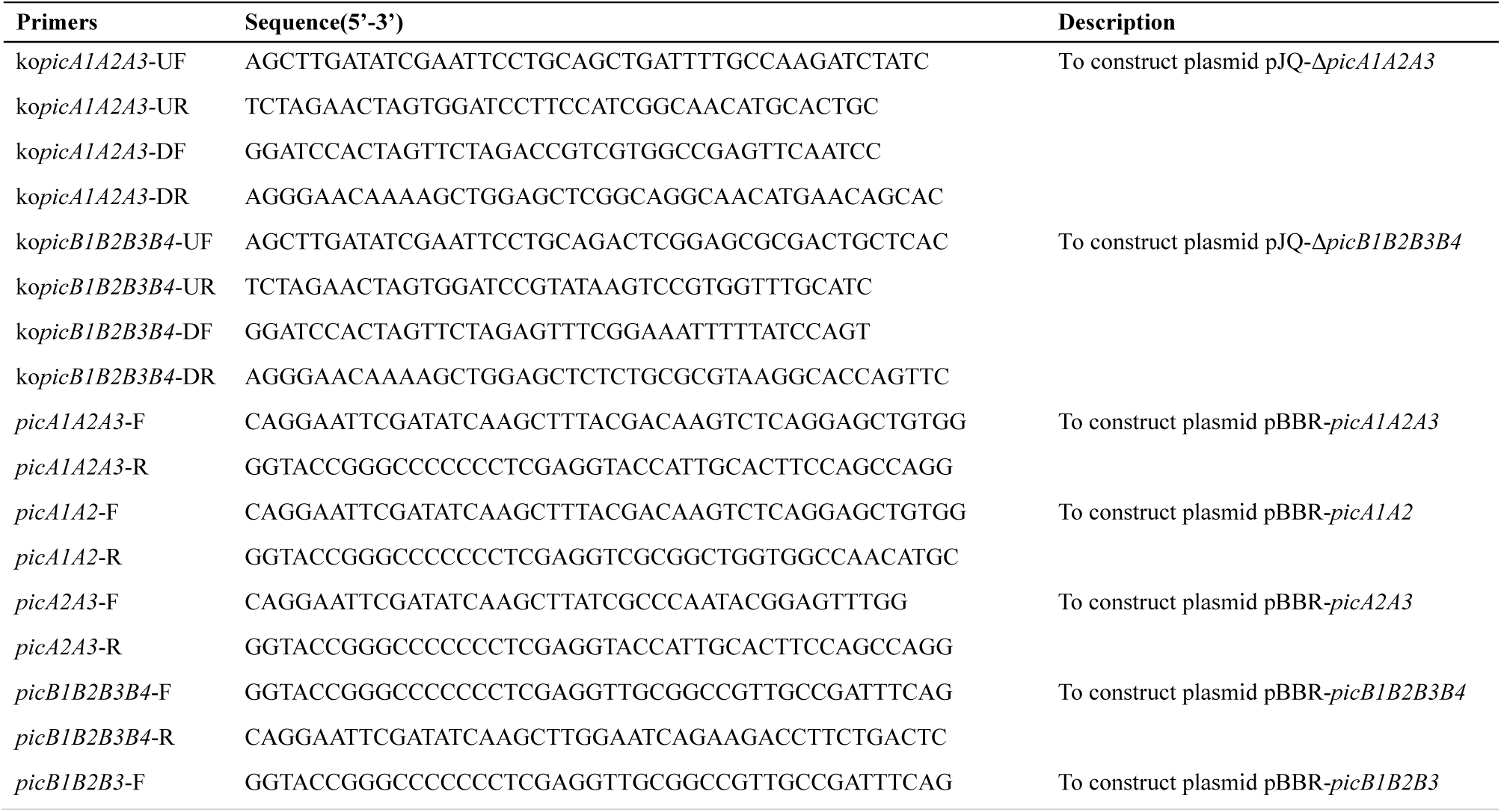
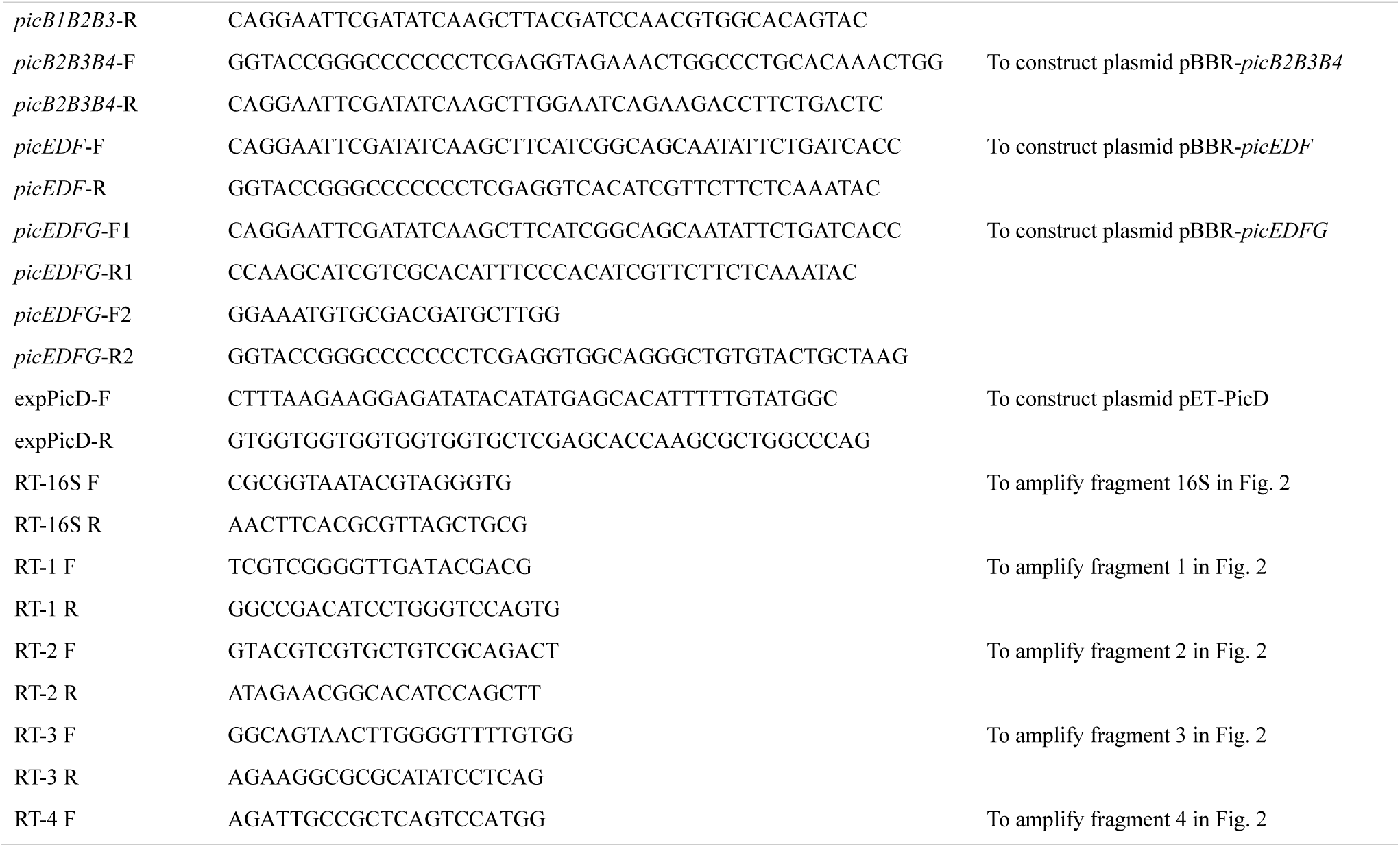
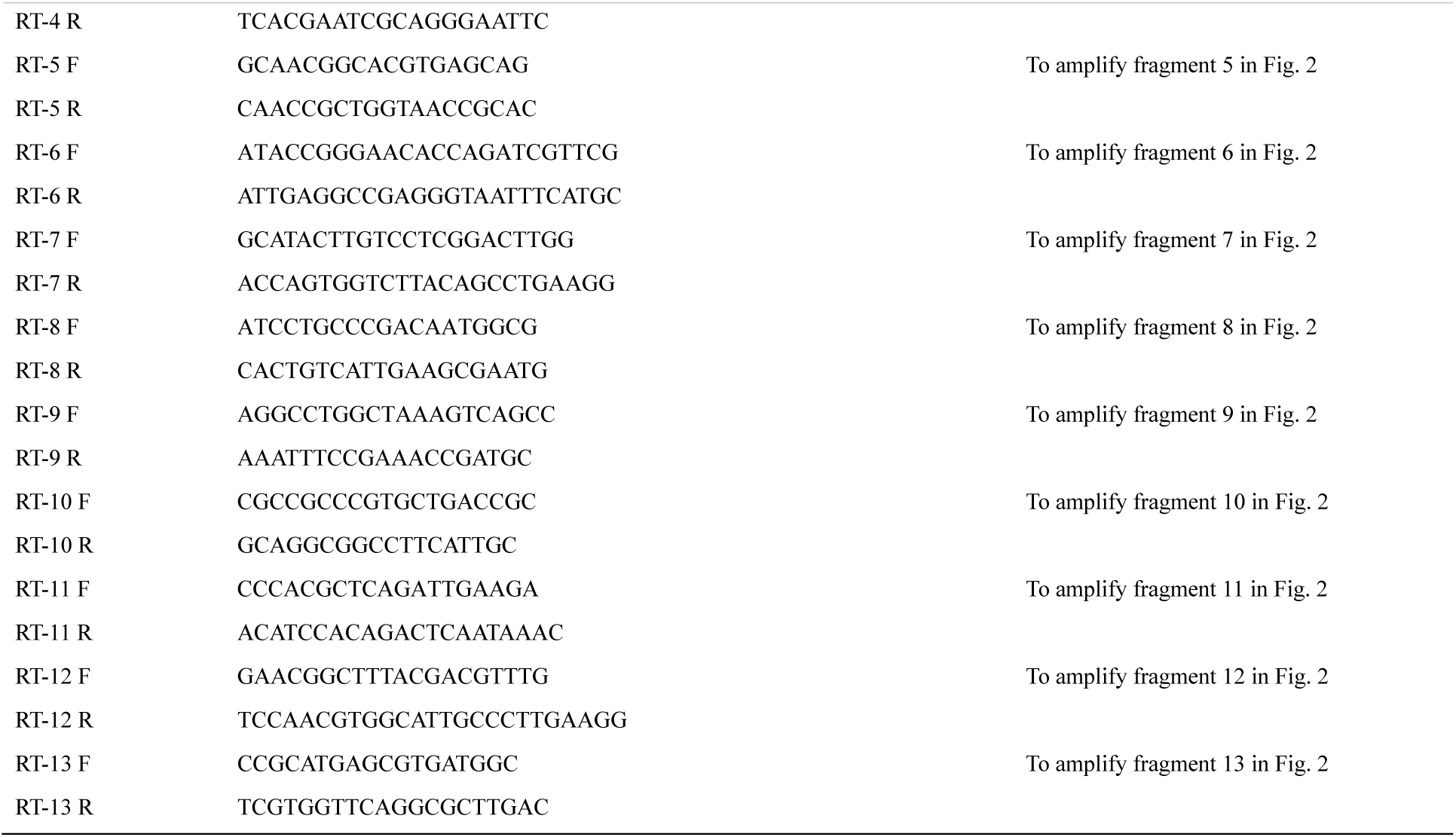
Primers used in this study.

### Construction of recombinant plasmids and heterologous expression

The genes from *A. faecalis* JQ135 were PCR amplified using corresponding primers (Table 3). The fusion of DNA fragments and cut plasmids were performed with the ClonExpress MultiS One Step Cloning Kit (Vazyme Biotech Co., Ltd, Nanjing, China).

The plasmid pBBR1MCS-5 (42) was used for heterologous expression in *P. putida* KT2440. PCR products and XhoI/HindIII-digested plasmid were ligated and the resulting recombinant plasmids were transferred into *P. putida* KT2440 by biparental mating using SM10_λpir_.

To test the components of PA dehydrogenase, pBBR1MCS-5-based plasmid clones containing different genes (*picA1A2A3, picA1A2*, and *picA2A3*) were transferred into *P. putida* KT2440, to yield recombinants KT/pBBR-*picA1A2A3*, KT/pBBR-*picA1A2*, and KT/pBBR-*picA2A3*. Similarly, to test the components of 6HPA monooxygenase, the strains KT/pBBR-*picB1B2B3B4*, KT/pBBR-*picB1B2B3*, and KT/pBBR-*picB2B3B4* were constructed.

For the over-expression of *picD* gene in *E. coli* BL21(DE3), the complete ORF without its corresponding stop codon was amplified and inserted into the NdeI/XhoI-digested plasmid pET29a(+), resulting in the plasmid pET-PicD. The inductuion and purification of 6×His-tagged PicD from *E. coli* BL21(DE3) (containing pET-PicD) were same with a previous report (25). The purified 6×His-tagged PicD were then analyzed by 12.5% SDS-PAGE. The protein concentrations were quantified using the Bradford method (43).

### Gene knockout and genetic complementation of *A. faecalis* JQ135

Deletion mutants of the *picA1A2A3* and *picB1B2B3B4* genes in *A. faecalis* JQ135 were constructed using a two-step homogenetic recombination method with the suicide plasmid pJQ200SK (44). Using the deletion of *picA1A2A3* as an example, two homologous recombination-directing sequences (500 to 1000 bp) were amplified using primers, ko*picA1A2A3*-UF/-UR and ko*picA1A2A3*-DF/-DR, respectively. The two PCR fragments were subsequently ligated into SacI/PstI-digested pJQ200SK yielding pJQ-Δ*picA1A2A3*. The pJQ-*ΔpicA1A2A3* plasmid was then introduced into *A. faecalis* JQ135 cells. The single-crossover mutants were screened on a LB plate containing Str and Gm. The gentamicin-resistant strains were then subjected to repeated cultivation in LB medium containing 10% sucrose and no gentamicin. The double-crossover mutants, which lost their vector backbone and were sensitive to gentamicin, were selected on LB Str plates. Deletion of the *picA1A2A3* genes was confirmed by PCR. This procedure resulted in construction of the deletion mutant strain JQ135Δ*picA1A2A3*.

Knockout mutants were complemented by the corresponding gene. For example, the intact *picA1A2A3* gene was amplified with primers *picA1A2A3*-F/-R, and then ligated with the XhoI/HindIII-digested pBBR1-MCS5, generating pBBR-*picA1A2A3*. The pBBR-*picA1A2A3* vector was transferred into the mutant strain JQ135Δ*picA1A2A3* to generate the complemented strain JQ135Δ*picA1A2A3*/pBBR-*picA1A2A3*.

### Preparation of resting cells and biodegradation assays

The *Alcaligenes, Pseudomonas*, and *Sphingomonas* strains and their derivates were grown in LB at 30 ºC. When the late exponential phase was reached, the cells were harvested by centrifugation at 6000 rpm for 10 min. The cells were washed twice by MSM and finally resuspended with MSM. The OD_600_ was adjusted to 2.0. These strains were named resting cells and were used for the biodegradation of PA or other related compounds.

### RT-PCR

Resting cells of *A. faecalis* JQ135 were inoculated into 20 mL MSM supplemented with 1 mM PA or citrate and cultured at 30ºC. The initial OD_600_ of the cultures was adjusted to 0.5. When half of the PA was degraded, the cells were harvested. Total RNA was isolated using an RNA isolation kit (TaKaRa) and reverse transcription-PCR (RT-PCR) was carried out with a PrimeScript RT reagent kit (TaKaRa). The primers used for RT-PCR are listed in Table 3. All samples were run in triplicate.

### Enzymatic assays

PA dehydrogenase (PicA) assays were performed using the cell-free extracts of KT/pBBR-*picA1A2A3*. Resting cells of KT/pBBR-*picA1A2A3* were added into MSM containing 1 mM citrate and 1 mM PA. Following incubation for 24 h, the cells were harvested by centrifugation at 6000 rpm for 10 min at 4 ºC. Cell pellets were then resuspended in 50 mM PBS (pH 7.0) and disrupted by sonication in an ice/water bath. Cell-free extracts were obtained by centrifuging the cell lysates at 16 000 g for 30 min at 4 ºC. The supernatant was later used for PA dehydrogenase assays. PicA activity was analyzed spectrophotometrically at 25 °C by the formation of 6HPA at 310 nm (ε=4.45 cm^−1^ mM^−1^) using a UV2450 spectrophotometer (Shimadzu) in 1 cm path length quartz cuvettes (23). The 1 mL reaction mixture contained 50 mM PBS (pH 7.0), 1 to 10 μg of total protein from cell extract, 1 mM PA, and 1 mM electron acceptor PMS and the reaction was started by adding PA. One unit of activity was defined as the amount of enzyme that catalyzed the formation of 1 μmol 6HPA in 1 min.

For the 6HPA monooxygenase (PicB) assays, the cell-free extracts of KT/pBBR-*picB1B2B3B4* were used. The preparation of cell-free extracts was the same as that of KT/pBBR-*picA1A2A3*. PicB activity was analyzed spectrophotometrically by looking for the formation of 3,6DHPA at 360 nm (ε=4.4cm^−1^ mM^−1^) (26). The 1 mL reaction mixture contained 50 mM PBS (pH 7.0), 1 to 10 μg of total protein from cell extract, 1 mM 6HPA, and 1 mM electron donor (NADH or others) and the reaction was started by adding 6HPA. One unit of activity was defined as the amount of enzyme that catalyzed the formation of 1 μmol 3,6DHPA in 1 min.

The 2,5-DHP dioxygenase (PicD) assays was performed similarly to those previously performed for NicX from *P. putida* KT2440 (28) and VppE in *Ochrobactrum* sp. strain SJY1 (33). A 1 mL reaction mixture contained 50 mM PBS (pH 7.0), 0.2 mM 2,5DHP, 0.1 μg purified PicD, and 1 mM Fe^2+^ and was incubated at 25 ºC. Activity was assayed spectrophotometrically by measuring the disappearance of 2,5DHP at 320 nm (ε=5.2 cm^−1^ mM^−1^). One unit of activity was defined as the amount of enzyme that catalyzed the consumption of 1 μmol 2,5DHP in 1 min.

## Analytical methods

The determination of PA and 6HPA, 3,6DHPA, and 2,5DHP were performed by Ultraviolet-visible spectroscopy and High performance liquid chromatography analysis were same with a previous study (26).

### Sequence data and bioinformatic analysis

The *pic* cluster sequence and the complete genome sequence of *Alcaligenes faecalis* JQ135 have been deposited in the GenBank/DDBJ/EMBL database under accession numbers <u>KY264362</u> and <u>CP021641</u>, respectively.

Comparisons of the Pic proteins were performed against the non-redundant protein sequences (nr) database using Blastp (protein-protein BLAST) programs on the NCBI website, employing an Expect (E)-value inclusion threshold of 10. The conserved protein domains of each gene were analyzed using the Conserved Domain Database (CDD; http://www.ncbi.nlm.nih.gov/Structure/cdd/wrpsb.cgi). The genome sequence accession numbers of other strains, and the corresponding locus tags of *pic* gene clusters, are listed in Table S1.

## Acknowledgments

This work was supported by the National Science and Technology Major Project (2018ZX0800907B-002), the National Natural Science Foundation of China (Nos.41630637, 31870092, 31770117, and 31600080). Key R&D Program Project in Jiangsu Province (BE2016374).

## References

1. Heyes MP, Eugene O, Saito K. 1997. Different kynurenine pathway enzymes limit quinolinic acid formation by various human cell types. Biochem J 326:351–356.

2. Bryleva EY, Brundin L. 2017. Kynurenine pathway metabolites and suicidality. Neuropharmacology 112:324–330.

3. Esquivel DG, Ramirez-Ortega D, Pineda B, Castro N, Rios C, de la Cruz VP. 2017. Kynurenine pathway metabolites and enzymes involved in redox reactions. Neuropharmacology 112:331–345.

4. Nishino SF, Spain JC. 1993. Degradation of nitrobenzene by a *Pseudomonas pseudoalcaligenes*. Appl Environ Microbiol 59:2520–2525.

5. Mehler AH. 1956. Formation of picolinic and quinolinic acids following enzymatic oxidation of 3-hydroxyanthranilic acid. J Biol Chem 218:241–254.

6. Asano Y, Yamamoto Y, Yamada H. 1994. Catechol 2, 3-dioxygenase-catalyzed synthesis of picolinic acids from catechols. Biosci Biotechnol Biochem 58:2054–2056.

7. Chirino B, Strahsburger E, Agullo L, Gonzalez M, Seeger M. 2013. Genomic and functional analyses of the 2-aminophenol catabolic pathway and partial conversion of its substrate into picolinic acid in *Burkholderia xenovorans* LB400. Plos One 8:e75746.

8. Abdullaev M, Klyuev M, Abdullaeva ZS, Kurbanov B, Idrisova A. 2008. Preparation of lidocaine, bipuvacaine, mepivacaine, trimecaine, and pyromecaine by reductive acylation on palladium catalysts. Pharm Chem J 42:357–359.

9. McCall PJ, Agin GL. 1985. Desorption kinetics of picloram as affected by residence time in the soil. Environ Toxicol Chem 4:37–44.

10. Broadhurst CL, Domenico P. 2006. Clinical studies on chromium picolinate supplementation in diabetes mellitus—a review. Diabetes Technol Therapeut 8:677–687.

11. Martin J, Wang ZQ, Zhang XH, Wachtel D, Volaufova J, Matthews DE, Cefalu WT. 2006. Chromium picolinate supplementation attenuates body weight gain and increases insulin sensitivity in subjects with type 2 diabetes. Diabetes Care 29:1826–1832.

12. Ogata S, Inoue K, Iwata K, Okumura K, Taguchi H. 2001. Apoptosis induced by picolinic acid-related compounds in HL-60 cells. Biosci Biotechnol Biochem 65:2337–2339.

13. Cioczek-Czuczwar A, Czuczwar P, Turski WA, Parada-Turska J. 2016. Influence of picolinic acid on seizure susceptibility in mice. Pharmacol Rep 69:77–80.

14. Prodinger J, Loacker LJ, Schmidt RL, Ratzinger F, Greiner G, Witzeneder N, Hoermann G, Jutz S, Pickl WF, Steinberger P. 2015. The tryptophan metabolite picolinic acid suppresses proliferation and metabolic activity of CD4+ T cells and inhibits c-Myc activation. J Leukocyte Biol 99:429–431.

15. Nakata HM, Halvorson HO. 1960. Biochemical changes occurring during growth and sporulation of *Bacillus cereus*. J Bacteriol 80:801–810.

16. Tamer Ö, Tamer SA, İdil Ö, Avcl D, Vural H, Atalay Y. 2017. Antimicrobial Activities, DNA interactions, spectroscopic (FT-IR and UV-Vis) characterizations, and DFT calculations for pyridine-2-carboxylic acid and its derivates. J Mol Struct 1152:399–408.

17. Suksrichavalit T, Prachayasittikul S, Nantasenamat C, Isarankura-Na-Ayudhya C, Prachayasittikul V. 2009. Copper complexes of pyridine derivatives with superoxide scavenging and antimicrobial activities. Euro J Med Chem 44:3259–3265.

18. Smythe GA, Poljak A, Bustamante S, Braga O, Maxwell A, Grant R, Sachdev P. 2003. ECNI GC-MS analysis of picolinic and quinolinic acids and their amides in human plasma, CSF, and brain tissue, p. 705–712, Developments in Tryptophan and Serotonin Metabolism. Springer.

19. Kiener A, Glockler R, Heinzmann K. 1993. Preparation of 6-oxo-1,6-dihydropyridine-2-carboxylic acid by microbial hydroxylation of pyridine-2-carboxylic acid. J Chem Soc Perk T:1201–1202.

20. Siegmund I, Koenig K, Andreesen JR. 1990. Molybdenum involvement in aerobic degradation of picolinic acid by *Arthrobacter picolinophilus*. FEMS Microbiol Lett 67:281–284.

21. Zheng C, Wang Q, Ning Y, Fan Y, Feng S, He C, Zhang TC, Shen Z. 2017. Isolation of a 2-picolinic acid-assimilating bacterium and its proposed degradation pathway. Bioresource Technol 245:681–688.

22. Zheng C, Zhou J, Wang J, Qu B, Wang J, Lu H, Zhao H. 2009. Aerobic degradation of 2-picolinic acid by a nitrobenzene-assimilating strain: *Streptomyces* sp. Z2. Bioresource Technol 100:2082–2084.

23. Orpin CG, Knight M, Evans WC. 1972. The bacterial oxidation of picolinamide, a photolytic product of Diquat. Biochem J 127:819–831.

24. Qiu J, Zhang J, Zhang Y, Wang Y, Tong L, Hong Q, He J. 2017. Biodegradation of picolinic acid by a newly isolated bacterium *Alcaligenes faecalis* strain JQ135. Curr Microbiol 74:508–514.

25. Qiu J, Liu B, Zhao L, Zhang Y, Cheng D, Yan X, Jiang J, Hong Q, He J. 2018. A novel degradation mechanism for pyridine derivatives in *Alcaligenes faecalis* JQ135. Appl Environ Microbiol 84:e00910–00918.

26. Qiu J, Zhang Y, Yao S, Ren H, Qian M, Hong Q, Lu Z, He J. 2019. Novel 3,6-dihydroxypicolinic acid decarboxylase mediated picolinic acid catabolism in *Alcaligenes faecalis* JQ135. J Bacteriol Accepted. (JB00665-18).

27. Dagley S, Johnson PA. 1963. Microbial oxidation of kynurenic, xanthurenic and picolinic acids. Biochim Biophys Acta 78:577–587.

28. Jimenez JI, Canales A, Jimenez-Barbero J, Ginalski K, Rychlewski L, Garcia JL, Diaz E. 2008. Deciphering the genetic determinants for aerobic nicotinic acid degradation: The nic cluster from *Pseudomonas putida* KT2440. Proc Natl Acad Sci USA 105:11329–11334.

29. Grether-Beck S, Igloi GL, Pust S, Schilz E, Decker K, Brandsch R. 1994. Structural analysis and molybdenum ? dependent expression of the pAO1 ? encoded nicotine dehydrogenase genes of *Arthrobacter nicotinovorans*. Mol Microbiol 13:929–936.

30. Chakraborty J, Ghosal D, Dutta A, Dutta TK. 2012. An insight into the origin and functional evolution of bacterial aromatic ring-hydroxylating oxygenases. J Biomol Struct Dyn 30:419–436.

31. Vaitekūnas J, Gasparavičiūtė R, Rutkienė R, Tauraitė D, Meškys R. 2016. A 2-hydroxypyridine catabolism pathway in *Rhodococcus rhodochrous* strain PY11. Appl Environ Microbiol 82:1264.

32. Tate RL, Ensign JC. 1974. Picolinic acid hydroxylase of *Arthrobacter picolinophilus*. Can J Microbiol 20:695–702.

33. Yu H, Tang H, Zhu X, Li Y, Xu P. 2015. Molecular mechanism of nicotine degradation by a newly isolated strain, *Ochrobactrum* sp. strain SJY1. Appl Environ Microbiol 81:272–281.

34. Petkevičius V, Vaitekūnas J, Stankevičiūtė J, Gasparavičiūtė R, Meškys R. 2018. Catabolism of 2-hydroxypyridine by *Burkholderia* sp. strain MAK1: a 2-hydroxypyridine 5-monooxygenase encoded by *hpdABCDE* catalyzes the first step of biodegradation. Appl Environ Microbiol 84:AEM.00387–00318.

35. Chen Q, Wang C-H, Deng S-K, Wu Y-D, Li Y, Yao L, Jiang J-D, Yan X, He J, Li S-P. 2014. Novel three-component Rieske non-heme iron oxygenase system catalyzing the *N*-dealkylation of chloroacetanilide herbicides in sphingomonads DC-6 and DC-2. Appl Environ Microbiol 80:5078–5085.

36. Strnad H, Ridl J, Paces J, Kolar M, Vlcek C, Paces V. 2011. Complete genome sequence of the haloaromatic acid-degrading bacterium *Achromobacter xylosoxidans* A8. J Bacteriol 193:791–792.

37. Mahan KM, Zheng H, Fida TT, Parry RJ, Graham DE, Spain JC. 2017. Iron-dependent enzyme catalyzes the initial step in biodegradation of *N*-nitroglycine by *Variovorax* sp. strain JS1663. Appl Environ Microbiol 83:e00457–00417.

38. Parkhill J, Sebaihia M, Preston A, Murphy LD, Thomson N, Harris DE, Holden MT, Churcher CM, Bentley SD, Mungall KL. 2003. Comparative analysis of the genome sequences of *Bordetella pertussis, Bordetella parapertussis* and *Bordetella bronchiseptica*. Nat Genet 3532.

39. Ju S, Lin J, Zheng J, Wang S, Zhou H, Sun M. 2016. *Alcaligenes faecalis* ZD02, a novel nematicidal bacterium with an extracellular serine protease virulence factor. Appl Environ Microbiol 82:2112–2120.

40. Vollmers J, Voget S, Dietrich S, Gollnow K, Smits M, Meyer K, Brinkhoff T, Simon M, Daniel R. 2013. Poles apart: Arctic and Antarctic *Octadecabacter* strains share high genome plasticity and a new type of xanthorhodopsin. Plos One 8:e63422.

41. Cubillas C, Miranda-Sánchez F, González-Sánchez A, Elizalde JP, Vinuesa P, Brom S, García-De LSA. 2017. A comprehensive phylogenetic analysis of copper transporting P1B ATPases from bacteria of the *Rhizobiales* order uncovers multiplicity, diversity and novel taxonomic subtypes. MicrobiologyOpen 6:e452.

42. Kovach M, Elzer P, Hill D, Robertson G, Farris M, Roop R, Peterson K. 1995. Four new derivatives of the broad-hostrange cloning vector pBBR1MCS, carrying different antibioticresistance cassettes. Gene 166:175–176.

43. Bradford MM. 1976. A rapid and sensitive method for the quantitation of microgram quantities of protein utilizing the principle of protein-dye binding. Anal Biochem 72:248–254.

44. Quandt J, Hynes M. 1993. Versatile suicide vectors which allow direct selection for gene replacement in gram-negative bacteria. Gene 127:15–21.

