## Supplemental figures for "Identification and characterization of a novel *pic* gene cluster responsible for picolinic acid degradation in *Alcaligenes faecalis* JQ135"

### Supplementary figures and tables

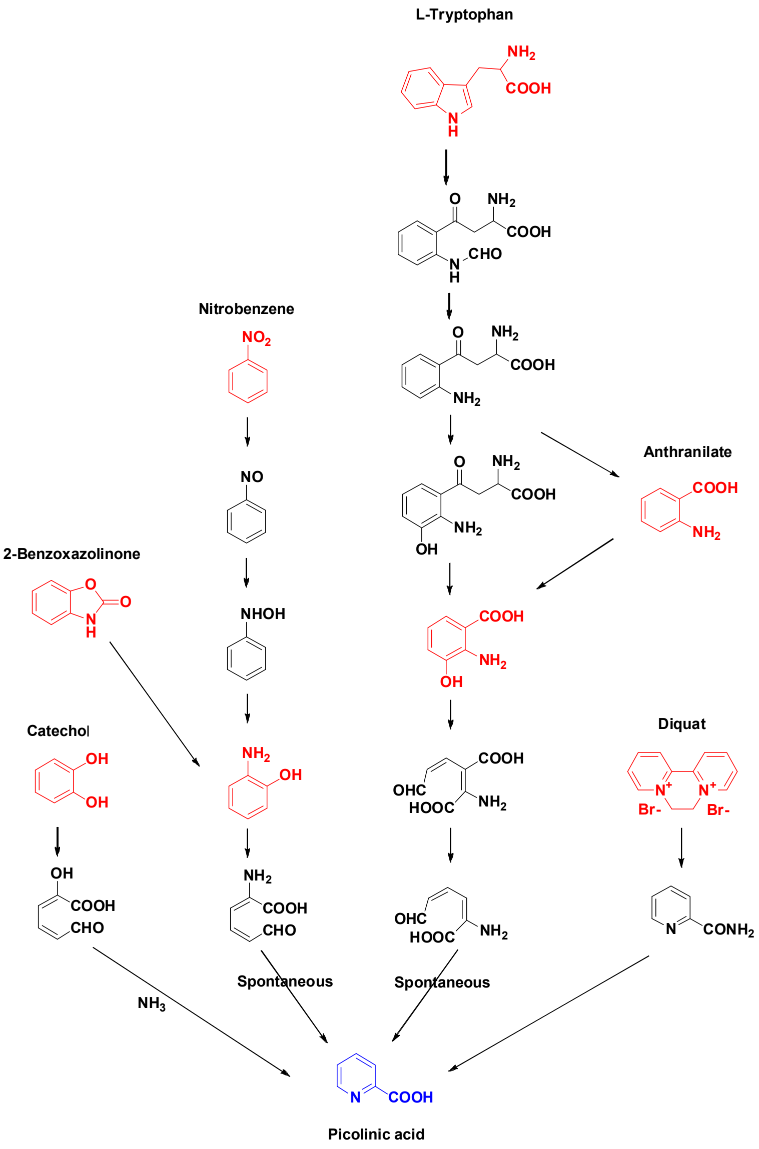


Fig. S1 Various sources of picolinic acid from the biological processes.

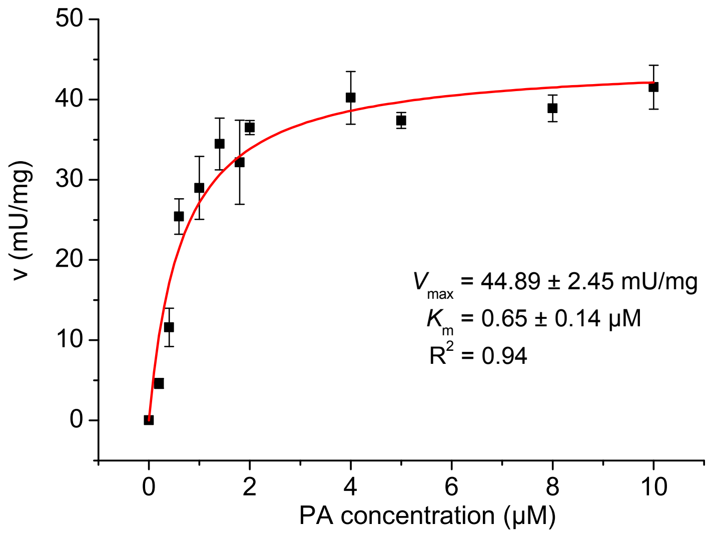


Fig. S2 The kinetic curves of PicA in cell-free extract of KT/pBBR-*picA1A2A3*. The Data were shown in means ± S.E.M.

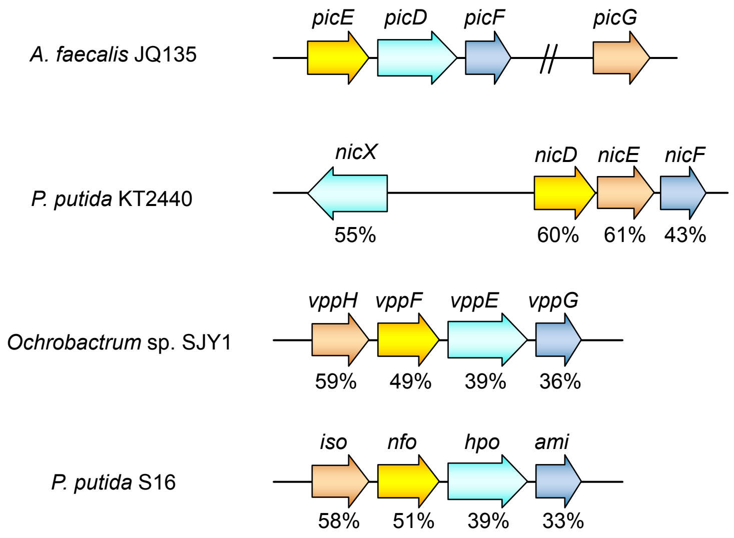


Fig. S3 Genetic organizations of the *picDEFG* genes with related genes involved in 2,5DHP degradation. The arrows indicate the size and direction of each gene. PicD (2,5DHP dioxygenase), PicE (*N*-formylmaleamate deformylase), PicF (maleamate amidohydrolase), and PicG (maleate *cis-trans* isomerase). Homologous genes are shown in the same color. Double vertical lines indicate discontinuous genes. Numbers below the arrows indicate the percent amino acid sequence identity with the ortholog *pic* gene product. *nicXDEF* (locus_tag PP_3945, PP_3943, PP_3942, and PP_3941) involved in nicotinate degradation from *P. putida* KT2440. *vppHFEG* (AIH15801, AIH15800, AIH15799, and AIH15798) involved in nicotine degradation from *Ochrobactrum* sp. SJY1. *iso*, *nfo*, *hpo* and *ami* (ADN26549, ADN26550, ADN26551, and ADN26552) involved in nicotine degradation from *P. putida* S16.

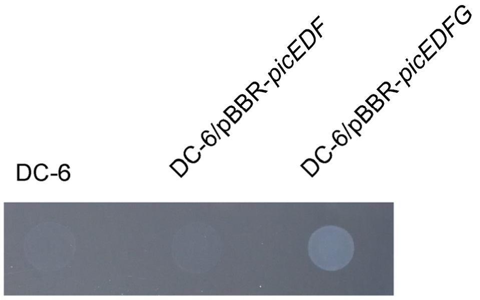


Fig. S4 Growth phenotype of the wild type *Sphingomonas wittichii* DC-6 and recombinant DC-6/pBBR-*picDEF* and DC-6/pBBR-*picDEFG*, on MSM plate with 1.0 mM 2,5-dihydroxypyridine as sole carbon source.

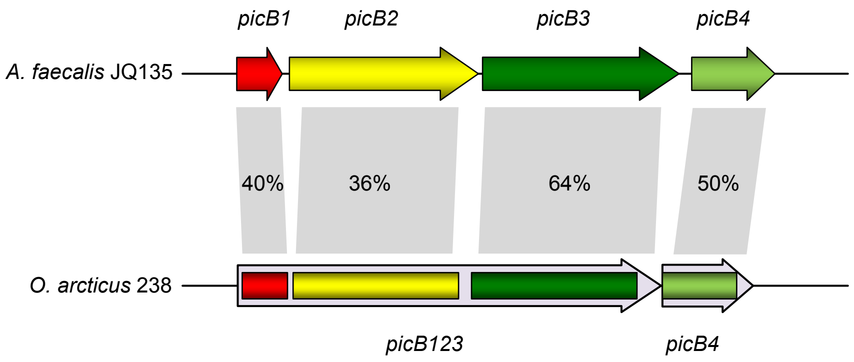


Fig. S5 The comparison of the PicB encoding genes of *A. faecalis* JQ135 and *O. arcticus* 238. Homologous genes/fragments are shown in the same color. Identities (%) of amino acid sequence of PicB components are indicated under the corresponding ORFs. In strain *O. arcticus* 238, the locus_tag of *picB123* and *picB4* are OA238_c08200 and OA238_c08210, respectively.
